# The defined TLR3 agonist, Nexavant, exhibits anti-cancer efficacy and potentiates anti-PD-1 antibody therapy by enhancing immune cell infiltration

**DOI:** 10.1101/2023.10.25.564097

**Authors:** Seung-Hwan Lee, Young-Ho Choi, Soon-Myung Kang, Min-Gyu Lee, Arnaud Debin, Eric Perouzel, Seung-Beom Hong, Dong-Ho Kim

## Abstract

Nexavant has been reported as an alternative to the TLR3 agonist of Poly(I:C) and its derivatives. The physicochemical properties, signaling pathways, anti-cancer effects, and mechanisms of Nexavant were investigated. Nexavant’s distinct nature, compared to Poly(I:C), was evident through precise quantification, thermostability, and resistance to RNase A. Unlike Poly (I: C) which activates TLR3, RIG-I and MDA5, Nexavant stimulates the signaling through TLR3 and RIG-I but not MDA5. Intratumoral Nexavant treatment led to a unique immune response compared to Poly(I:C), immune cell infiltration, and suppression of tumor growth in various animal cancer models. Nexavant therapy outperformed anti-PD-1 antibody treatment in all tested models and showed a synergistic effect in combinational therapy, especially in well-defined cold tumor models. The effect was similar to Nivolumab in a humanized mouse model. Intranasal instillation of Nexavant recruited immune cells (NK, CD4+ T, CD8+ T) to the lungs, suppressing lung metastasis and improving animal survival. Our study highlighted Nexavant’s defined nature for clinical use, unique signaling pathways, and its potential as a standalone anti-cancer agent or in combination with anti-PD-1 antibody.

**Simple Summary:** Nexavant, a newly reported TLR3 agonist, has advantages over Poly(I:C) in quality control and pre-clinical efficacy. Here, we further investigated Nexavant’s physicochemical properties, downstream signaling pathways, anti-cancer efficacy, and mechanism of action. Nexavant was homogenous in solution, less sensitive to RNase A, and showed thermostability compared with Poly(I:C). Unlike Poly(I:C), the TLR3, RIGI, and MDA5 activator, Nexavant only activated TLR3 and RIG-I but not MDA5. Administration of Nexavant either by intratumoral or intranasal route suppressed tumor growth in various cancer models. Combination therapy with anti-PD-1 antibody exhibited synergistic tumor growth inhibition than the respective monotherapies. This study demonstrated that Nexavant could be more suitable for clinical use over Poly(I:C) and applied as an anti-cancer agent in the presence or absence of an anti-PD-1 antibody.

## 1. Introduction

Since the approval of Yervoy, a CTLA-4-targeting antibody, for treating melanoma in 2011, immune checkpoint inhibitors (ICIs), such as anti-PD-1 and PD-L1 antibodies, have been widely used to treat cancer1, 2. However, apart from a small percentage of patients with high response rates to ICIs (approximately 10-35%), many patients still have no or low response rates, and some patients develop resistance to ICIs, resulting in cancer recurrence or accelerated cancer growth3, 4. The main reason for the development of resistance to immuno-oncology drugs, including ICIs, is the increased secretion of immunosuppressive cytokines and angiogenic factors from immunosuppressive cells and the inhibition of immune cell infiltration into cancer cells (cold tumor, immuno-cold). Transforming a “cold tumor” with inhibited immune cell infiltration into a “hot tumor” with more immune cells can maximize the efficacy of immuno-oncology drugs and significantly improve treatment response rates^5^.

To this end, research and clinical trials are underway to deliver pathogen-associated molecular patterns (PAMPs), such as Poly(I:C), to tumors^6-9^. PAMPs are recognized by pattern recognition receptors (PRRs), a protein found in the cell membrane or cytoplasm, to induce an innate immune response in the host. This allows a variety of immune cells to infiltrate the tumor and transform it into a hot tumor. In situ vaccines, injected directly into the tumor, can induce an immune response specific to an individual’s cancer by using a variety of antigens present in cancer. Since it can utilize various antigens in cancer cells, it has the advantage of efficiently inducing immune responses to multiple cancer antigens and overcoming the heterogeneity of cancer cells.

Toll-like receptor 3 (TLR3) and retinoic acid-induced gene-I (RIG-I)-like receptors (RLR) are well-known PRRs that recognize dsRNA and are highly expressed on innate immune cells such as DCs and macrophages, etc^9-11^. These receptors serve as the first line defense system of the immune system, allowing the host to recognize invading foreign pathogens with dsRNA. Their activation leads to the production of type I interferon (IFN), proinflammatory cytokines, and chemokines through activation of interferon-stimulated gene (ISG) signaling, inducing infiltration of various immune cells and a broad immune response^12, 13^.

A well-known representative dsRNA is Polyinosinic: Polycytidylic acid, abbreviated as Poly(I:C). It can strongly induce the secretion of IFN-β and inflammatory cytokines by activating TLR3, RIG-I, and MDA5 and can cause the infiltration of various immune cells^9, 11, 14, 15^. In this regard, It is attracting attention as vaccine adjuvants and in situ vaccines. Poly(I:C) is synthesized poly-inosine and poly-cytidine conjugates with dsRNAs’ structure but have unordered complementarities, resulting in high heterogeneity^16, 17^. The nature of Poly(I:C) significantly challenges stability, quality control, and monitoring in vivo PK. Various attempts have been made to overcome these obstacles to improve the stability. The diverse formulation was tried by adding additives to Poly(I:C) to produce Poly-ICLC, Poly-IC12U, PICKA, and BO-112^9, 16^. However, all of them use Poly(I:C) as the centerpiece, so the limitations of Poly(I:C) still exist. Another solution is the length-determined candidates by chemical syntheses, such as RGC100, ARNAX, and TL-532^16, 18^. As shown by other studies^19^, these molecules are not for the most optimized length for activation of TLR3. Nevertheless, the potent efficacy of Poly(I:C) has encouraged researchers to conduct various clinical trials. In particular, poly-ICLC (Hiltonol) is undergoing dozens of clinical trials as a vaccine adjuvant and in situ vaccine for various solid cancer patients^16, 20^.

We recently presented Nexavant as a well-defined TLR3 agonist^17^. Nexavant offers several advantages over Poly(I:C) regarding quality control and clinical applicability. Nexavant comprises a 424 bp dsRNA core with five nt single-stranded overhangs at both 3’ ends, giving it a defined molecular weight of 275 kDa. Importantly, Nexavant demonstrated stability under accelerated conditions (25°C) for six months and a safety profile in several pre-clinical animal studies17. Unlike Poly(I:C), Nexavant can be produced up to 99% purity with a scale-up and cost-effective synthesis. Moreover, it provides accurate pharmacokinetic and pharmacodynamic data, offering significant regulatory advantages^17^.

Intramuscular administration of Nexavant to mouse systems notably increases the migration of dendritic cells (DCs), macrophages, and neutrophils to the draining lymph nodes. Nexavant also triggers the upregulation of MHC-II, CD40, CD80, and CD86 in DCs, influencing DC maturation and activation. Nexavant holds promise as a vaccine adjuvant, as it efficiently stimulates dendritic cells, pivotal cells that impact vaccine efficacy. The activation of dendritic cells by Nexavant enhances their antigen-presenting capacity and cytokine secretion. It may ultimately increase cancer antigen-specific T-cell responses and alterations in the cancer microenvironment, leading to enhanced anti-cancer effects.

In this study, we investigated the physicochemical properties, downstream signaling pathways, and anti-cancer efficacy of Nexavant compared with those of Poly(I:C). Simple methods can precisely monitor the absolute amount and quality of Nexavant. In addition, the anti-cancer effect of Nexavant and Poly (I:C) by intratumoral or intranasal administration was compared in various animal models. Considering the enhanced anti-cancer effect, signaling pathway, and possible molecules involved, Nexavant was proposed as an alternative tool for anticancer therapy of cold tumors. Furthermore, the efficacy of anti-PD1 antibody therapy was synergistically enhanced when combined with intratumoral administration of Nexavant, especially in the cold tumor models. Nexavant treatment led to immune cell infiltration and suppressed tumor growth by a unique innate immune response.

## 2. Materials and Methods

### 2.1. Animals

Female C57bl/6 or Balb/c mice at 6-8 weeks of age mice were purchased from Samtake Bio Korea (Kyounggi, Korea), and hPD-1-knockin C57bl/6 or Balb/c mice were purchased from GH Bio (Daejeon, Korea). The mice were maintained at the NA Vaccine Institute (NAVI) animal facility (Seoul, Korea), fed a sterile, commercial mouse diet, and provided with water *ad libitum*. The experimental protocols used in this study were reviewed and approved by the NAVI’s Ethics Committee and Institutional Animal Care and Use Committee (IACUC).

### 2.2. Cell lines

B16F10 murine melanoma cell line was purchased from Korean Cell Line Bank (Seoul, Korea), and CT26 murine colon cancer cell line, LL/2 murine lung cancer cell line, and 4T-1 murine breast cancer cell line were purchased from ATCC. B16F10, LL/2, and EMT6 cells were maintained in DMEM (Welgene, Korea), and CT26 and 4T-1 cells were maintained in RPMI (Welgen). All DMEM and RPMI media were supplemented with 10% fetal bovine serum (FBS) (Welgene) and 1% penicillin/streptomycin (Gibco) and incubated at 37°C and 5% CO^2^.

HEK-Dual RNA 4KO hTLR3, HEK-Dual RNA 4KO hRIG-I, and HEK-Dual RNA 4KO MDA5 cell lines were developed by InvivoGen (CA, USA). They are derived from the human embryonic kidney 293 (HEK293)-Dual cell line harboring the stable integration of two inducible reporter genes for SEAP (secreted embryonic alkaline phosphatase) and Lucia luciferase. As a result, these cells allow the simultaneous study of the NF-κB pathway by monitoring the activity of SEAP and the IRF (interferon regulatory factor) pathway by assessing the activity of a secreted luciferase.

Briefly, HEK-Dual RNA 4KO cells were obtained using standard genome editing procedures to suppress the gene expression of TLR3 together with RIG-I, MDA5, and PKR. These cells’ complete absence of IRF responses upon stimulation with dsRNAs has been functionally validated. HEK-Dual RNA 4KO cells were engineered to express only one dsRNA sensor stably, using TLR3-, RIG-I, or MDA5-encoding plasmids. The SEAP, Lucia, and PRR expression was maintained by growing the cells in media containing Blasticidin (ant-bl-05, InvivoGen), Zeocin (ant-zn-05, InvivoGen), and Puromycin (ant-pr-1, InvivoGen). Cell lines were routinely checked for mycoplasma contamination using PlasmoTest (rep-pt-1, InvivoGen).

### 2.3. Reagents and antibodies

Nexavant was produced by In vitro transcription following the procedures described in the previous report17. Poly(I:C) HMW (tlrl-pic for in vitro assay, vac-pic for in vivo assay), Poly(I:C) LMW (Cat. #.: tlrl-picw), the lipid-based transfection reagent LyoVec (Cat. #.: lyec-12), the luciferase detection reagent QUANTI-Luc 4 Lucia/Gaussia (Cat. #.: rep-qlc4lg1) and the SEAP detection reagent QUANTI-Blue 4 (Cat. #.: rep-qbs) were purchased from InvivoGen. Cell staining antibodies for the flow cytometry analysis, including PE/Cy7-conjugated anti-mouse CD11b mAb (Cat. #.: 101215), APC-conjugated anti-mouse CD11c mAb (Cat. #.: 117310), FITC-conjugated anti-mouse CD19 mAb (Cat.#.: 115506), FITC-conjugated anti-mouse CD335 mAb (Cat. #.: 137606), PE/Cy5-conjugated anti-mouse CD3 mAb (Cat. #.: 100309), PE/Cy5-conjugated anti-mouse CD45 mAb (Cat. #.: 103110), PerCP/Cyanine5.5-conjugated anti-mouse CD45 mAb (Cat. #.: 103132), PE-conjugated anti-mouse CD80 mAb (Cat. #.: 104708), PE/Cy5-conjugated anti-mouse CD86 mAb (Cat. #.: 105016), APC/Cyanine7-conjugated anti-mouse CD8 mAb (Cat. #.: 100714), FITC-conjugated anti-mouse CD8 mAb (Cat. #.: 100705), PE-conjugated anti-mouse F4/80 mAb (Cat. #.: 123110), AF700-conjugated anti-mouse I-A/I-E mAb (Cat. #.: 107621), PerCP/Cy5.5-conjugated anti-mouse Ly6G mAb (Cat. #.: 127615) and 7-AAD Viability Staining Solution (Cat. #.: 420404) were purchased from BioLegend (CA, USA).

### 2.4. Treatment of Nexavant and Poly(I:C)

When Nexavant and Poly(I:C) were administered to mice, two administration methods were used depending on the experiment. For intratumoral administration, 50 ul of Nexavant and Poly(I:C) were slowly injected into the center of the tumor using an Ultra-Fine II syringe (BD, NJ, USA). For nasal delivery to the lungs, mice were respiratory anesthetized with isoflurane, and 30 ul of Nexavant and Poly(I:C) was slowly instilled dropwise through the nostrils.

### 2.5. ISG signaling induction

In a 96-well plate, 5 x 104 HEK-Dual RNA 4KO hTLR3, HEK-Dual RNA 4KO hRIG-I, or HEK-Dual RNA 4KO MDA5 cells were cultured overnight with increasing concentrations of Poly(I:C) HMW (Cat. #.: tlrl-pic, InvivoGen), Poly(I:C) LMW (Cat. #.: tlrl-picw, InvivoGen), or Nexavant, either naked (10 µg/ml to 10 ng/ml) or pre-complexed (1 µ/ml to 1 ng/ml) with the LyoVec transfection reagent according to manufacturer instructions. Recombinant human IFN-β (1000 U/ml to 1 U/ml, ThermoFisher) was used as positive controls for IRF activation. According to manufacturer instructions, the ISG response was measured by monitoring the Lucia luciferase activity in the culture supernatants using QUANTI-Luc 4 Lucia/Gaussia.

### 2.6. Tumor models

5 x 10^5^ cells of B16F10 or LL/2 cells were subcutaneously inoculated into the flank of C57bl/6 mice, and CT26 cells were inoculated into Balb/c mice. 4T-1, EO771, or EMT6 cells were inoculated into the mammary fat pad of Balb/c mice. Nexavant or Poly(I:C) HMW (vac-pic, InvivoGen) was injected into the tumors of the mice, and an anti-mouse PD-1 antibody (Cat. #: BE0146, BioXCell, NH, USA), was injected intraperitoneally at the same time. Tumor size was measured with a caliper three to four times a week, and tumor volume was calculated using the equation (width x length x height) x 3.14 / 6. Mice were euthanized when exhibiting signs of poor health or when the tumor volume exceeded about 2,000 mm3. To establish the lung metastasis model, 5 x 10^5^ cells of B16F10 were injected through the tail vein of C57bl/6 mice. As the experimental schedule, the lung metastasis mice were sacrificed, and the lungs were harvested. The gross photograph of a lung with tumor burden was taken with a digital camera (Sony α5100), and the tumor nodules of the lungs were counted.

### 2.7. Analysis of immune cells from inguinal lymph nodes and tumors

The inguinal lymph nodes were collected at the indicated times from the mice injected with Nexavant and crushed into a 70 μm strainer. Then, the cells of the lymph nodes were dissociated with collagenase type IV (Worthington Biochemical Co., NJ, USA) for 20 min at 37°C, washed with PBS including 1% FBS and 0.1% sodium azide (Sigma Aldrich, MO, USA), and subjected to flow cytometry. The tumors were collected on day 13 from the B16F10 tumor-bearing mice and crushed into a 70 μm strainer. Then, the tumor cells were dissociated with collagenase type IV and I (Worthington Biochemical Co.) for 20 min at 37°C and washed with PBS containing 1% FBS and 0.1% sodium azide (Sigma Aldrich). Cells were stained with antibodies, as described above. Unstained samples were used as negative controls. Stained cells were analyzed using a NovoCyte flow cytometer (Agilent Technologies, CA, USA).

### 2.8. Enzyme-linked immunosorbent assay (ELISA)

Blood samples were collected from the mice treated with Nexavant or the mice bearing tumors. Serum was separated from the blood samples after centrifugation at 13000 rpm for 10 min. The serum’s level of IFN-β, IL-6, and IL-12 was quantified with ELISA kits according to the manufacturer’s instructions (Invitrogen, CA, USA).

### 2.9. Statistical analysis

Data were analyzed using Prism V6 (GraphPad, CA, USA) and represented as mean values ± standard error of the mean (SEM). One-way ANOVA with Dunnett’s test was performed to compare more than two groups. *p<0.05 was considered statistically significant.

## 3. Results

### 3.1. Physicochemical advantage of Nexavant as a homogeneous TLR3 agonist

We compared the physicochemical properties of Nexavant and Poly(I:C). First, the feasibility of quantitation using a simple UV-absorbance spectrometry was tested. Poly(I:C) was notably evident with relatively low accuracy (coefficient of variation, 29.48%). The peak absorbance was observed between 250-270 nm. The absorption spectrum of Poly(I:C) exhibited two maxima at λ = 248 nm and λ = 269 nm (Figure 1A). These characteristics are likely attributed to the low uniformity within the solution and the heterogeneous structure of Poly(I:C), which consists of a mixture of single- and double-stranded RNA with varying chain lengths. Nexavant measurements were highly accurate (coefficient of variation, 1.56%), and their absorbance profiles resembled those of typical nucleic acids, showing a peak at 260 nm. Unlike Poly(I:C), the amount of Nexavant can be monitored by simple UV spectrometry at any time. By adopting simple spectrometry and agarose gel analyses, the quality and quantity of Nexavant can be measured conveniently.

**Figure 1.**
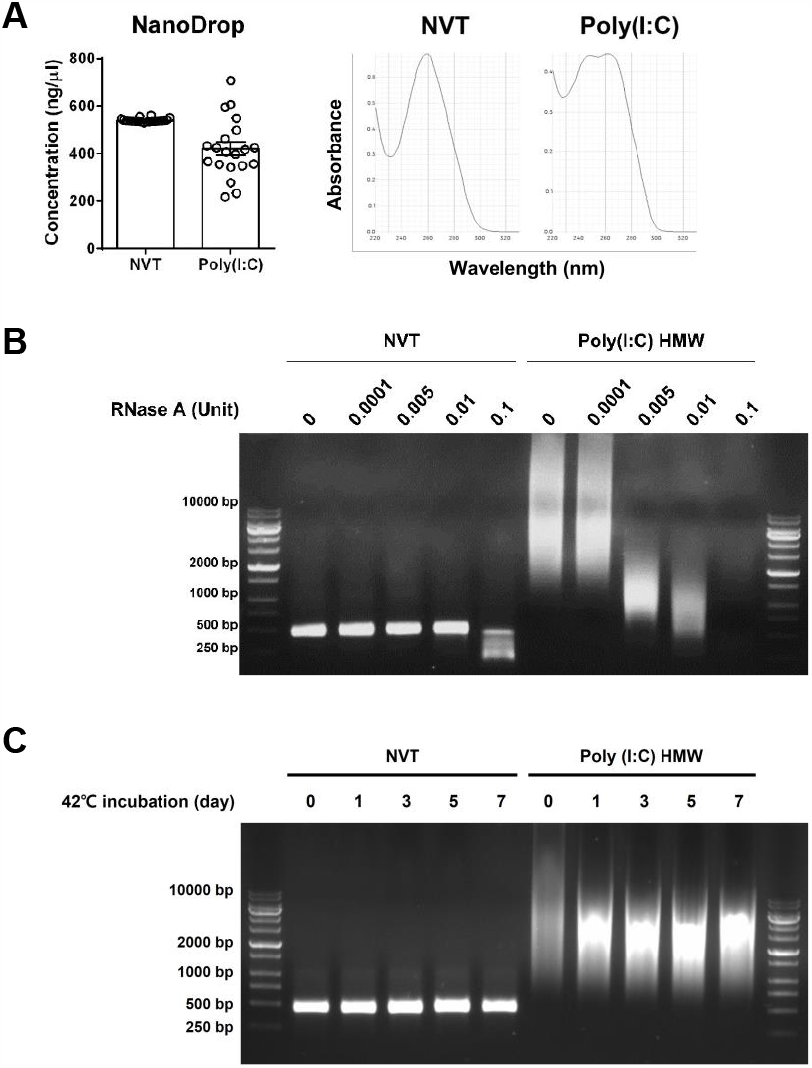
Physicochemical characteristics of Nexavant (NVT) and Poly(I:C) as demonstrated by UV spectrometric analysis, stability to RNase A, and thermostability (A) Accuracy of quantification by UV-spectrometry (A260 nm) and profiles of UV-absorbance spectra (NanoDrop, *n*=20). (B) Susceptibility to various concentrations of RNase A at 37oC for 10 min. (C) Susceptibility to different incubation times at 42°C.

We observed that Nexavant displayed lower sensitivity to RNase A than Poly(I:C) (Figure 1B). While Poly(I:C) underwent degradation at low doses of RNase A concentrations of 0.005 U/mL, Nexavant remained resistant to RNase A even at higher concentrations. The susceptibility of Poly(I:C) to RNase A could be attributed to the presence of single-stranded RNA generated during the manufacturing process due to incomplete hybridization between poly-I and poly-C chains (Figure 1B). In contrast, Nexavant’s reduced sensitivity to RNase A can be attributed to its dsRNA structure with complete complementary hybridization. When each molecule was stored at 42°C for thermostability assay, Poly(I:C) exhibited structural changes or material alterations within a day. Nexavant remained stable after a week (Figure 1C) and six months (data not shown). These findings highlight that Nexavant is a well-defined TLR3 agonist in terms of its physical and structural properties. It can be consistently produced, exhibits low sensitivity to RNase A, and demonstrates long-term stability at non-freezing temperatures.

### 3.2. Similar but distinct TLR3 pathway activation of Nexavant

MDA5 detects long-duplex RNAs in the genome of double-stranded RNA (dsRNA) viruses or dsRNA replication intermediates of viruses21, 22. By contrast, RIG-I detects the five triphosphate group (5′ppp) and the blunt end of short dsRNAs or single-stranded RNA (ssRNA) hairpins23, 24. To determine the cellular action mechanisms of Nexavant, we applied HEK-293 dual reporter model cell lines for interferon stimulating signaling, which express either TLR3, RIG-I, or MDA5 genes ectopically on the basis of genetic deficiency of four different endogenous dsRNA sensor genes (i.e., TLR3, RIG-I, MDA5, and PKR). RIG-I, MDA5, and TLR3 are strongly activated when Poly(I:C) is transfected into cells using cationic lipids (LyoVec). Nexavant turned out to be a decent TLR3 and RIG-I activator regardless of the presence of the lipid, while it did not induce MDA5 activation even with LyoVec. Consistent with other studies showing the MDA5 activation and dsRNA length dependency21, Nexavant is a novel TLR3 agonist that only activates RIG-I but not MDA5. Overall, Nexavant can be defined as a distinct TLR3 agonist with moderate potency and a unique activator of the molecular cascade system compared with Poly(I:C).

### 3.3. Intratumoral delivery of Nexnt induces the recruitment of various immune cells into the tumor

The immune cell profile in the inguinal lymph nodes and cytokine levels in the blood were monitored over 48 hours after a single subcutaneous injection of Nexavant on the flank of the healthy mouse (Figure 3A). Most of the analyzed immune cells, including CD4+ and CD8+ T cells, B cells, neutrophils, NK cells, and macrophages, were recruited to lymph nodes, although the time to peak and the kinetics were cell type-specific. The total number of dendritic cells in the lymph node remained constant. However, the expression of DC activation markers such as CD80 and CD86 increased over time, peaked at 24 hr, and declined afterward. The serum levels of IFN-β and proinflammatory cytokines such as IL-6 and IL-12, typical products of ISG signaling, were induced at 4 hr after injection (Figure 3B). While the levels of both IFN-β and IL-6 declined to undetectable levels by 12 hr, the levels of IL-12 remained longer and declined to baseline by 48 hr.

**Figure 2.**
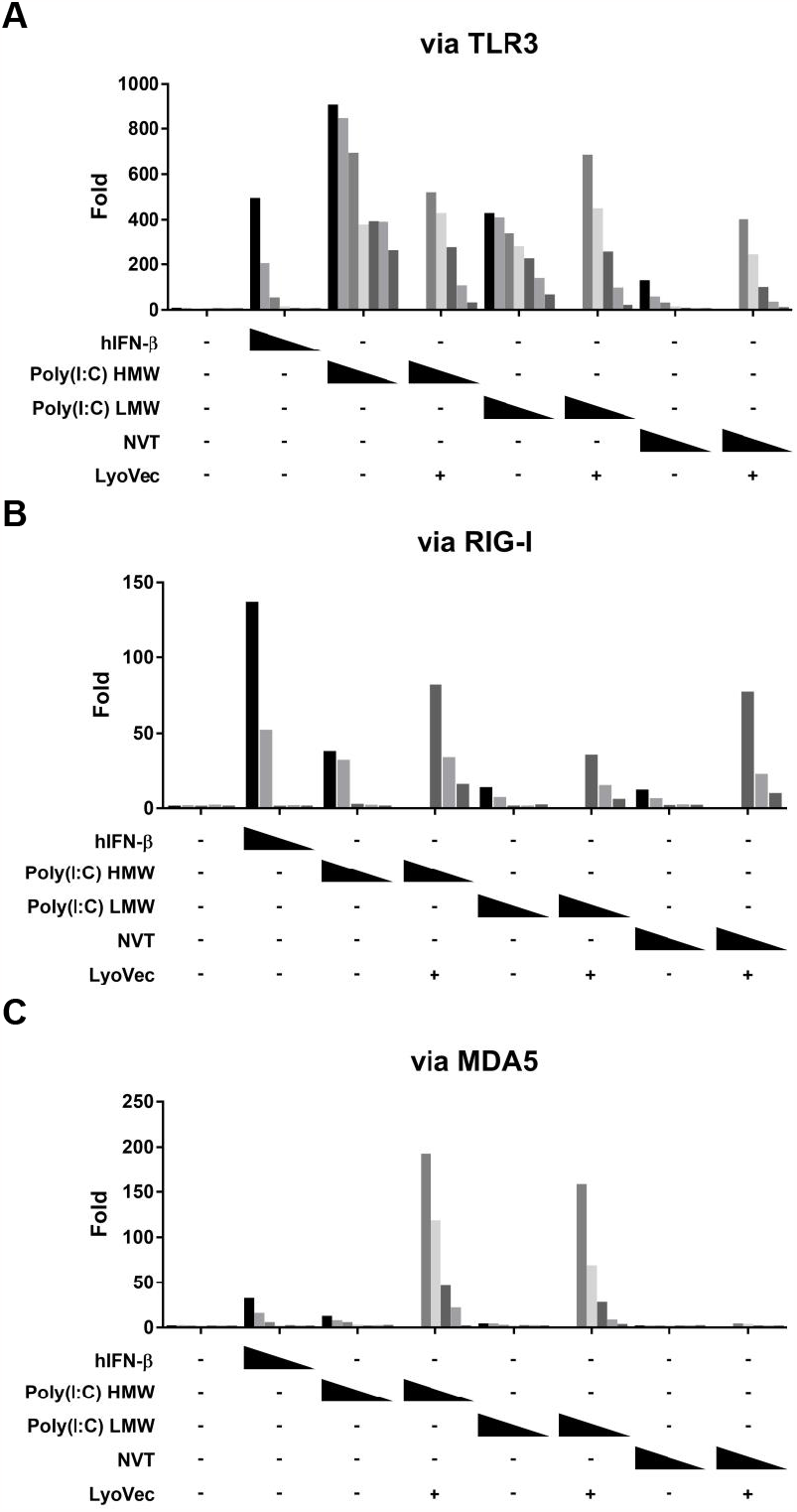
The naked and cationic-lipid-complexed Nexavant regulates the downstream signaling pathways. The three model cell lines expressing either TLR3, RIG-I, or MDA5 genes were stimulated with naked or cationic-lipid complexed Nexavant to examine its mechanism of action. Responsive reporter systems measured dose-dependent TLR3, RIG-I, and MDA5 signaling induction. As controls, hIFN-β, hIL-1β, Poly(I:C) HMW, and Poly(I:C) LMW were used. The fold inductions for RLU and absorbance compared with the non-treated groups were calculated. hIFN-β, Poly(I:C) HMW, Poly(I:C) LMW, and Nexavant were treated as follows. hIFN-β: 1000, 300, 100, 30, 10, 3, 1 Unit/ml; Poly(I:C) HMW, Poly(I:C) LMW or Nexavant: 10000, 3000, 1000, 300, 100, 30, 10 ng/ml; Poly(I:C) HMW, Poly(I:C) LMW or Nexavant (with LyoVec): 1000, 300, 100, 30, 10, 3, 1 ng/ml. Data are representative of two independent experiments.

**Figure 3.**
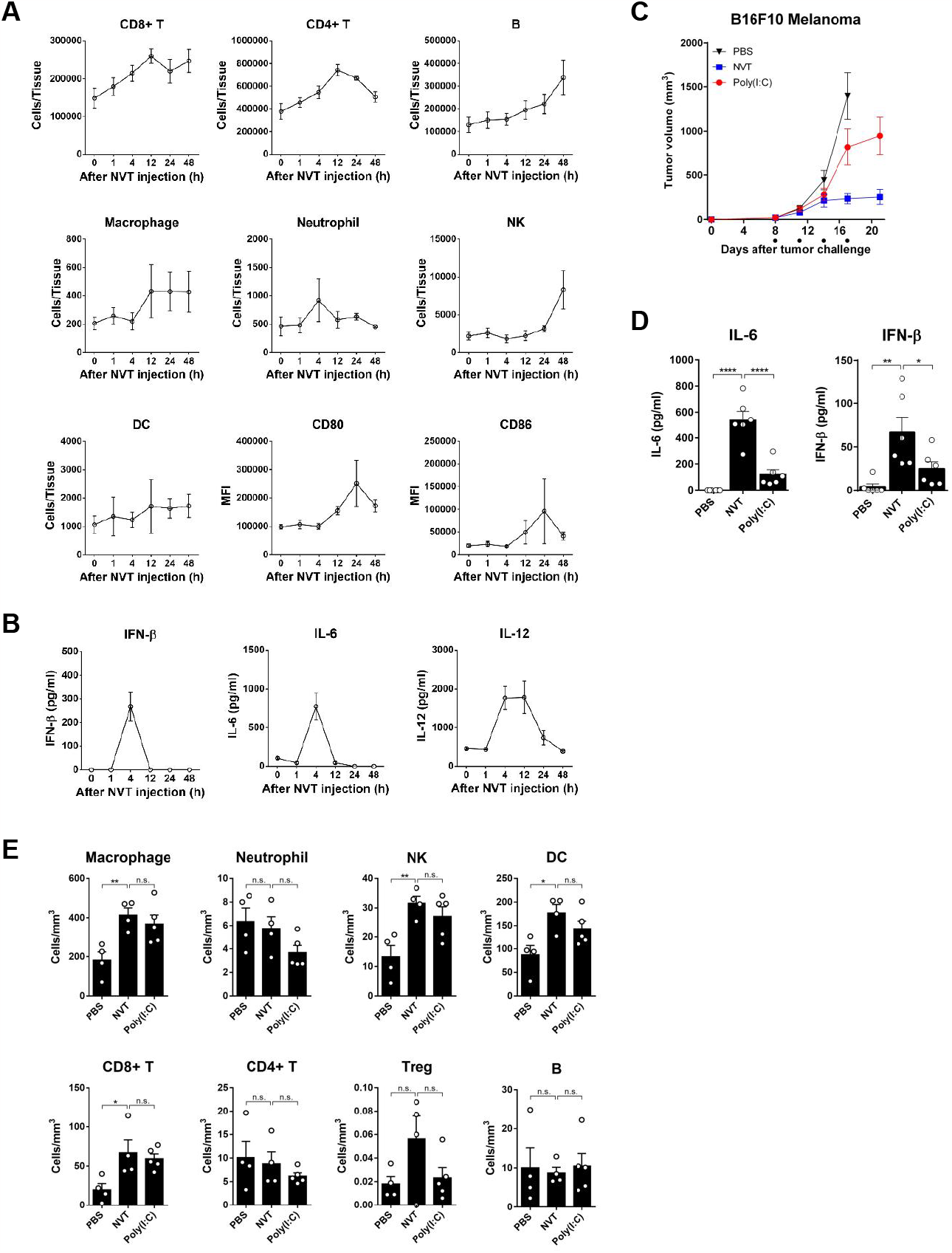
Nexavant recruits diverse immune cells to the draining lymph nodes near the injection sites. C57bl/6 mice were injected subcutaneously with 25 ug of Nexavant. At 1, 4, 12, 24, and 48 hours post-injection, the mice were monitored for (A) immune cell profile and DC activation in the inguinal lymph nodes and (B) IFN-β, IL-6 and IL-12 cytokine levels in the blood. C57BL/6 mice inoculated subcutaneously with 5 x 105 cells of B16F10 cells were treated with 100 ug Nexavant or Poly(I:C) on 8, 11, 14, and 17 days post-inoculation, and (C) Tumor growth was measured. (D) Blood levels of IL-6 and IFN-β cytokines were measured 4 hours after the first injection of Nexavant or Poly(I:C). (E) The immune cell profile of mouse tumors was then analyzed on day 15.

To examine Nexavant’s anti-cancer efficacy compared to Poly(I:C), each was administered to mouse tumors in the melanoma model, respectively (Figure 3C). While both suppressed tumor growth, Nexavant suppressed better than Poly(I:C) at the same dose (100 ug/animal; Tumor growth inhibition (TGI)% for Nexavant 83.18%, TGI% for Poly(I:C) 41.21%). To understand why Nexavant exhibits better anti-cancer efficacy, we examined the levels of cytokine induction in serum and immune cell infiltration into tumors after both treatments. The levels of IL-6 and IFN-β in the blood were higher in the group treated with Nexavant than the correspondingPoly(I:C) groups (Figure 3D). Immune cells such as macrophages, NK, DC, and CD8+ T cells within the tumors were significantly increased in both groups (Figure 3E). However, no statistical differences were found between the two groups.

The results demonstrated that Nexavant in the tumor tissue induces infiltration of diverse immune cells similar to Poly(I:C). Based on the better anti-cancer efficacy of Nexavant than Poly(I:C), the enhanced cytokine secretion, such as IFN-β and IL-6, may induce a unique tumor microenvironment for improved anticancer effect.

### 3.4. Intratumoral delivery of Nexavant shows anti-tumor effects in various tumor models

To investigate whether Nexavant shows anti-cancer effects beyond mouse melanoma, tumor growth inhibition by Nexavant was tested in various mouse cancer models (Figure 4A). Nexavant suppressed tumor growth by 61.5, 76.15, and 58.04% in LL/2 lung cancer, CT26 colon cancer, and 4T-1 breast cancer models, respectively. This indicates that Nexavant has broad anti-cancer effects on a variety of cancers. The dose-response and abscopal effect of Nexavant was examined in a mouse melanoma model (B16F10) (Figure 4B). Intratumoral administration of Nexavant at doses as low as 25 ug and as high as 150 ug resulted in significant tumor growth inhibition in both treated (72.93-96.49%) and untreated distant tumors (TGI% 62.09-74.73%). However, the dose-response was not evident, especially in the case of the ascobal effect.

**Figure 4.**
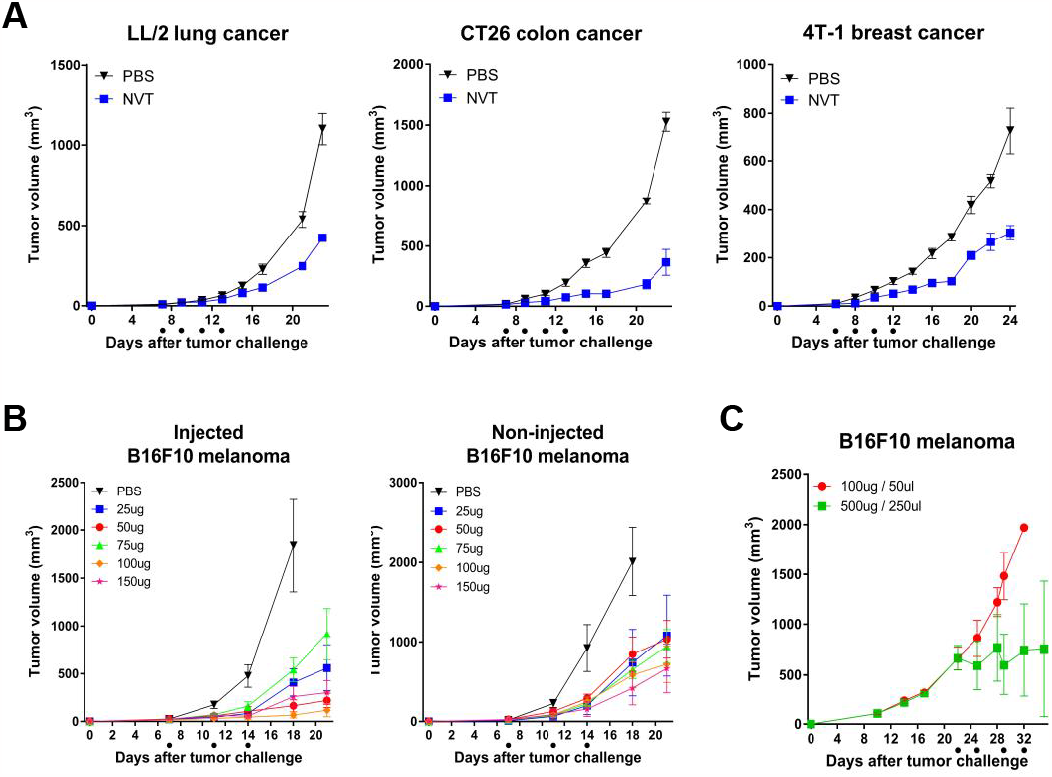
Intratumoral administration of Nexavant suppressed tumor growth in various syngeneic models. (A) C57bl/6 mice inoculated subcutaneously with 5 x 105 cells of LL/2, CT26, or 4T-1 cells were treated with Nexavant (25 ug) (on 7, 9, 11, and 13 days post-inoculation, LL/2 model and CT26 model; on 6, 8, 10 and 12 days post-inoculation, 4T-1 model). (B) C57BL/6 mice were inoculated subcutaneously with 5 x 105 B16F10 cells at both flanks. Nexavant (25, 50, 75, 100, 150 ugs) was injected into the right flank tumor 7, 11, and 14 days post-inoculation. (C) C57BL/6 mice inoculated subcutaneously with 5 x 105 cells of B16F10 cells were treated with Nexavant (100 ug and 500 ug) on 22, 25, 29, and 32 days post-inoculation.

To investigate if Nexavant is effective in the late stage of the tumor, it was administered 22 days after tumor cell injection (mean tumor volume 750 mm3) twice weekly. While 100 ug/animal was ineffective, 500 ug/animal effectively controlled tumor growth (Figure 4D). In this late-stage tumor-bearing model, tumor growth was suppressed when a higher dose was administered. This showed that the dose of Nexavant for intratumoral application should be varied depending on the volume and stage of the tumor. The presented data demonstrated that Nexavant has anti-cancer activity against a wide range of cancers, and the effective dose can be affected by the tumor status, such as tumor size and stage.

### 3.5. The anti-tumor effect of anti-PD-1 antibody therapy can be improved by delivering Nexavant intratumorally

We used mouse melanoma and TNBC models, known as cold tumors25, 26, to determine whether intratumoral administration of Nexavant combined with anti-PD-1 antibodies improves the anti-cancer effect. In the mouse melanoma model, the anti-PD-1 antibody and Nexavant monotherapy groups showed a TGI effect of 35.88 and 78.33%, respectively. The combination therapy group showed a TGI effect of 96.19% (Figure 5A). The treated animals harboring tumors were rechallenged with mouse melanoma cells on the flank of the opposite side on day 24 and monitored for tumor growth. No tumor growth from rechallenge was observed in the groups pre-treated with anti-PD-1 and combination therapy.

**Figure 5.**
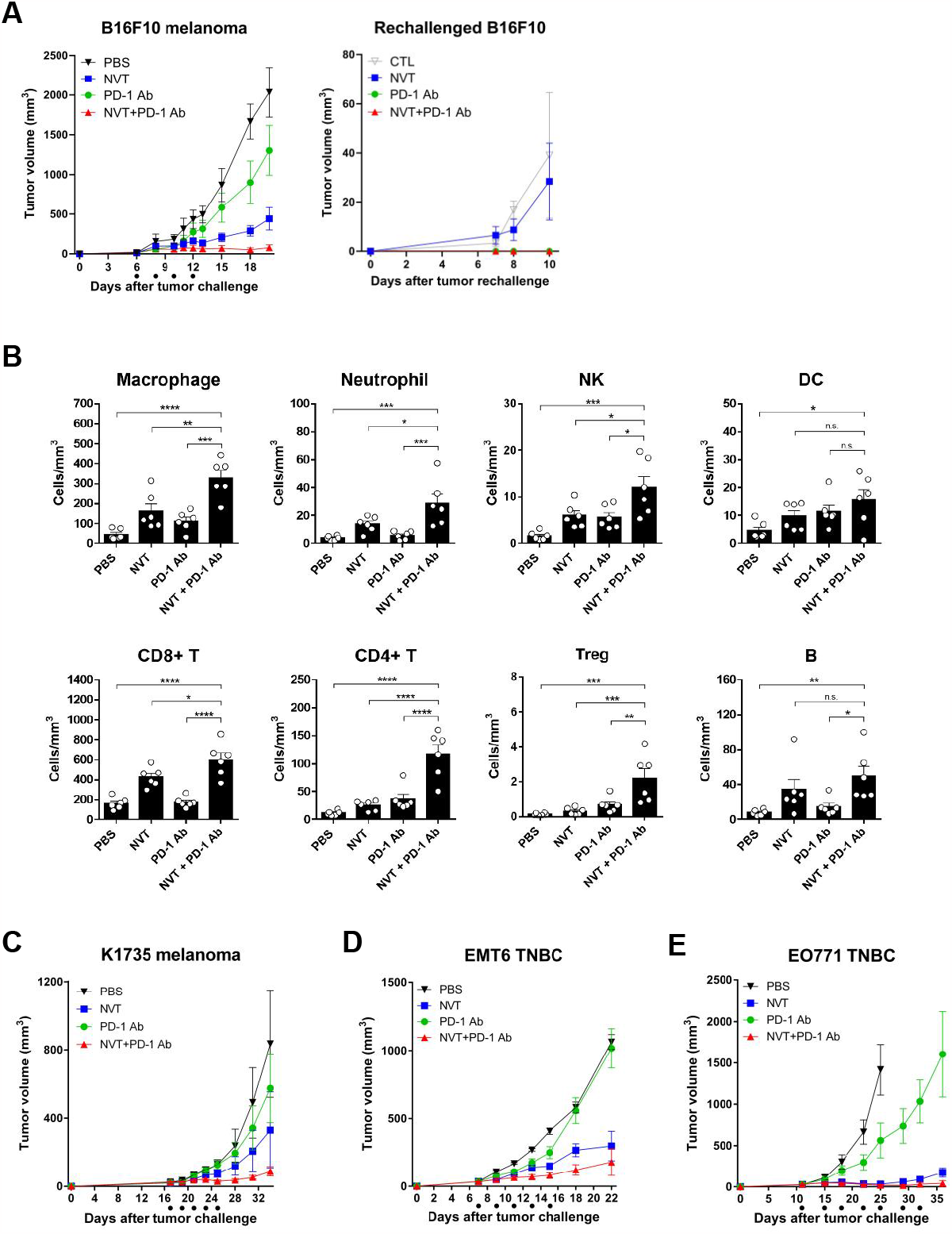
A combination of anti-PD-1 antibody therapy with Nexavant shows an improved anti-tumor effect. (A) C57bl/6 mice were inoculated subcutaneously with 5 x 105 cells of B16F10 tumor cells at the right flank and were treated with 25 ug Nexavant (intratumoral) and 100 ug anti-PD-1 antibody (intraperitoneal) on 6, 8, 10 and 12 days post-inoculation. 5 x 105 B16F10 tumor cells were inoculated again at the left flank on day 24 after the first challenge (corresponds to day 0 of the rechallenge). The volume of injected and non-injected tumors was measured at the indicated time points. (B) The immune cell profiles in the B16F10 tumor on day 25 were assessed by flow cytometry. (C) C57bl/6 mice were subcutaneously inoculated with 5 x 105 cells of K1735 tumor cells at the right flank and were immunized with Nexavant and anti-PD-1 antibody on 17, 19, 21, 23, and 25 days post-inoculation. (D) Balb/c mice were inoculated with 5 x 105 cells of EMT6 tumor cells into the mammary fat pad and then immunized with Nexavant and anti-PD-1 antibody on 7, 9, 11, 13, and 15 days post-inoculation. (E) C57bl/6 mice were inoculated with 5 x 105 cells of EO771 tumor cells into the mammary fat pad and then immunized with Nexavant and anti-PD-1 antibody on 11, 15, 18, 22, 25, 29, and 32 days post-inoculation. Tumor volumes were measured at indicated time points.

Intratumoral immune cell analysis demonstrated that combining anti-PD-1 antibody and Nexavant significantly increased NK cells, CD4, and CD8 T cells compared with the respective monotherapies (Figure 5B). The additive/synergistic effect was demonstrated not only in B16F10 melanoma but also in K1735 melanoma (Figure 5C). In the K1735 melanoma model, the tumor growth inhibition effects of Nexavant and anti-PD-1 antibody were 60.42% and 31.27%, respectively, and the combination significantly improved the inhibition effect to 89.46%. We also tested the effectiveness of the combination therapy in triple-negative breast cancer (TNBC), a well-known model of cold tumors (Figure 5D, E). The combination significantly improved the tumor inhibition effect in both models compared to the respective monotherapies. Although the degree of anti-cancer effect of Nexavant was different depending on the type of the tumor, it was confirmed that the combination therapy significantly improved the anti-cancer effect of anti-PD-1 antibody. The synergistic effect of dual therapy was evident in the melanoma and TNBC models, good examples of cold tumors.

### 3.6. Intratumoral administration of Nexavant also has a synergistic effect with Nivolumab, a human anti-PD-1 antibody

More human clinical relevant mode was pursued to investigate the efficacy of Nexavant and a human anti-PD-1 antibody, Nivolumab. We used a genetically engineered mouse expressing a chimeric PD-1 with the extracellular domain of human PD-1 and the transmembrane and cytoplasmic domains of mouse PD-1. Similar to the previous study using an anti-PD-1 antibody of murine origin, the combination of Nexavant and Nivolumab was more effective than the respective monotherapies in both B16F10 melanoma and EMT6 TNBC models (Figure 6A and 6B, respectively). While Nivolumab alone showed low tumor growth inhibition (49.85%), similar to the anti-PD-1 antibody of murine origin, the combination therapy dramatically improved tumor growth inhibition and survival. The combinational effect in the EMT6 TNBC model was impressive, with an 83% complete response. This further supports that the combined use of Nexavant can improve the anti-cancer efficacy of anti-PD-1 antibody therapy, especially in cold tumor models.

**Figure 6.**
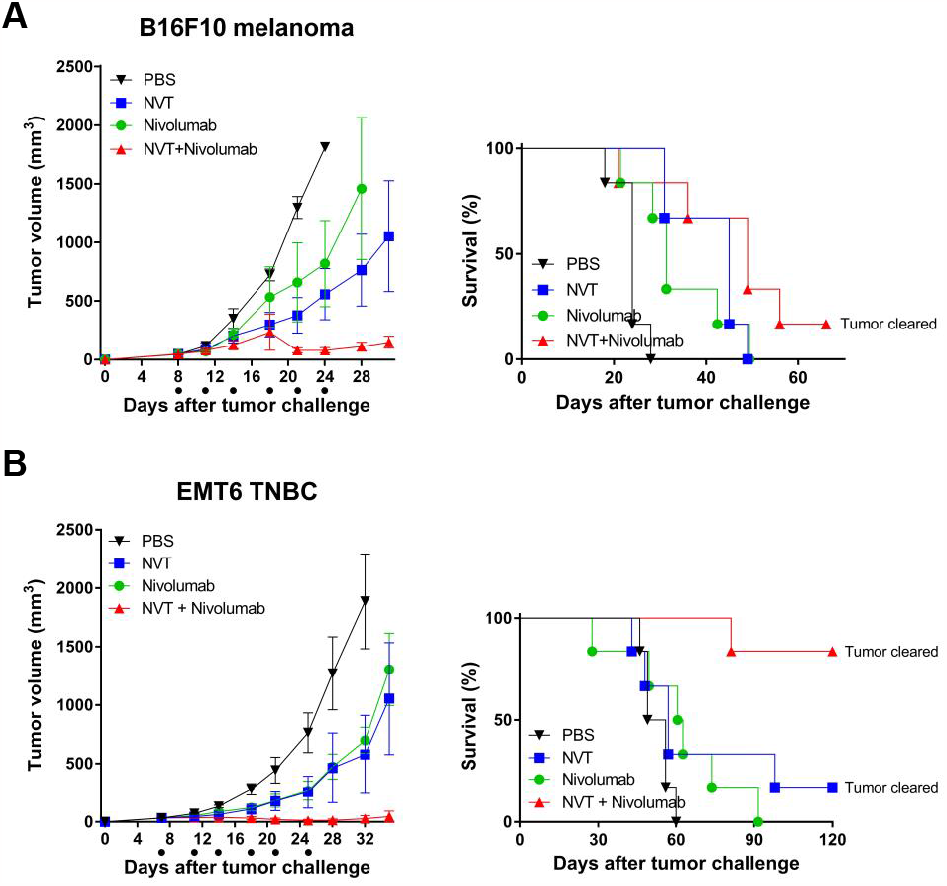
Improved anti-cancer efficacy by the combined treatment of Nexavant with Nivolumab (a human anti-PD-1 antibody) was tested in the transgenic mice expressing a chimeric PD-1 with the extracellular domain of human PD-1 and the transmembrane and cytoplasmic domains of mouse PD-1 compared with the respective monotherapies. C57BL/6 and Balb/c mice were inoculated with 5 x 105 cells of B16F10 melanoma cells and EMT6 TNBC cells, respectively (Day 0). (A, B) Tumor growth inhibition and improved survival by mono-vs. combination therapy in (A) B16F10 melanoma and (B) EMT6 breast cancer models. B16F10-bearing mice were treated with Nexavant and anti-PD-1 antibody on 8, 11, 14, 18, 21, and 24 days post-inoculation, and EMT6-bearing mice were treated on 7, 11, 14, 18, 21, and 25 days post-inoculation. Tumor volumes and survivals were measured at indicated time points.

### 3.7. Intranasal administration of Nexavant has a potent anti-tumor effect on the lung metastasis model

Poly(I:C) has been shown to have an anticancer effect on the lung cancer model through lung delivery27. To broaden the application of Nexavant, we tested whether Nexavant also has an anti-cancer effect when delivered to the lung metastasis model. We first examined how the lung’s immune cell profile changes when Nexavant is administered to the lung (Figure 7A). Flow cytometric analysis of the bronchoalveolar lavage fluid (BALF) cells harvested 24 hours after Nexavant administration demonstrated that NK and CD8+ T cells were significantly increased. No significant changes were found for macrophages, neutrophils, and CD4+ T cells. We tested whether Nexavant could inhibit lung metastasis when administered to the lung. In the B16F10 lung metastasis model, Nexavant or Poly(I:C) were instilled on days 8 and 14, and lung metastasis nodules on day 18. Compared to Poly(I:C), the lungs of Nexavant-treated mice showed a reduced number and size of tumor nodules (Figure 7B) and improved survival compared to the Poly(I:C) group (Figure 7C).

To investigate whether the stage of tumor development affects the anti-cancer effects of Nexavant, it was instilled every other day starting on 2 (early treatment group), 8 (intermediate treatment group), and 14 (late treatment group) days after cancer cell injection through tail vein and the survival rates of the animals were assessed (Figure 7D). Nexavant prolonged the survival of the animals when treated earlier (days 2 and 8) but did not when treated late (day 14). On day 15, the number of tumor nodules in the lungs of mice in the early treatment group, which responded best to Nexavant, was reduced about 7-fold compared with the controls (Figure 7E). Given the synergistic effects of intratumorally-administered Nexavant and anti-PD-1 antibody, we tested whether intranasally-administered Nexavant also enhances the efficacy of anti-PD-1 antibody treatment. While Nexavant monotherapy showed strong tumor growth suppression compared with anti-PD-1 antibody monotherapy, combination therapy did not exhibit better tumor growth suppression compared with the monotherapy. These results suggest that Nexavant could be very effective in some contexts and may not require combination therapy with an anti-PD-1 antibody.

## 4. Discussion

Poly(I:C) and its derivatives have been used as TLR3 agonists for more than a decade for diverse studies and clinical developments9, 16, 18. Despite tremendous studies, the clinical products based on them are minimal. In contrast to efficacy, they are hard to control the quality of homogeneity and quantitation in vitro and in vivo. Since no method is defined for the quantitation and qualification of Poly(I:C) and its derivates, the quality assurance of the sample is not feasible. Because Poly(I:C) is formed by incomplete annealing of poly-inosine and poly-cytosine chains, it is highly heterogeneous in length and structure, resulting in a heterogeneous macromolecule. Due to this heterogeneity, there is sizeable batch-to-batch variability in average length, composition, and in vivo stability, which may result in variability in efficacy.

Nexavant, a newly reported TLR3 agonist, is a dsRNA with a defined length and structure with a high uniformity17. Nexavant is a perfectly matched dsRNA with a specified length of 424 bp, which is expected to be more stable in vitro and in vivo than Poly(I:C). It has been shown that Nexavant is more resistant to RNase A digestion and shows thermostability at 42°C (Figure 1B and C). The homogeneous nature of Nexavant allows for better production quality control, which is expected to result in more consistent efficacy. Most of all, the amount and quality of Nexavant can be monitored by a simple method of UV absorbance or agarose gel electrophoresis. Since Nexavant can be stored at 42°C for up to 6 months (data not shown), logistics and storage of the reagent are convenient and energy and cost-efficient.

When RNase digestion converts the HW Poly(I:C) to a shorter form, the TLR3 agonist maintains RIG-I activation but loses the MDA5 activation21. Nexavant is shorter than Poly (I: C), and its activation was limited to RIG-I but not MDA5 (Figure 2). Based on this study, Nexavant should be classified as a defined TLR3 agonist for exclusive activators of RIG-I without MDA5. Compared with Nexavant, naked Poly(I:C) stimulated the dsRNA sensors with more potency. The differences in structure, length, and homogeneity likely contribute to the differences in the kinetics of cellular delivery and the activation of downstream signaling pathways. We previously reported that administration of Nexavant to mice increased mRNA levels of TLR3, RIG-I, and MDA5, suggesting that Nexavant may act as a dsRNA ligand similar to Poly(I:C)17. It is speculated that the induction of MDA5 signaling observed in vivo is due to a positive feedback effect on the activation of TLR3 and RIG-I signaling. In contrast to Poly(I:C) HMW, Nexavant induced relatively low TLR3 and RIG-I ISG signaling activation. Interestingly, it showed more significant tumor growth inhibition than Poly(I:C) in mouse tumor models (Figure 3C). This is likely due to differences in the in vivo stability and mode of action of Nexavant and Poly(I:C). More potent induction of IFN-β and IL-6 by Nexavant indicates the molecular difference, which should be defined further (Figure 3D).

Intratumoral administration of Nexavant induced recruitment of various immune cells to the administration site (Figure 3E). Nexavant may stimulate antigen-presenting cells (APCs) to capture diverse cancer antigens and migrate to draining lymph nodes to present cancer antigen information to T cells. Cancer antigen-specific T cells circulate throughout the body via blood or lymphatic vessels. They infiltrate to cancerous tissue where the cancer antigen is present and engage with cancer cells bearing the antigen. Eventually, the antigen-specific T cells become activated and eliminate the cancer cells. Due to their systemic circulation, these T cells have the potential to target and eradicate not only the primary cancer site but also metastatic cancers, a phenomenon known as the abscopal effect. When Nexavant is injected into a tumor, the immune cell infiltration pattern resembles Poly(I:C) (Figure 3E, and citation28). Based on the difference in efficacy and the level of IFN-β and IL-6, the potent anticancer effect of Nexavant may need to be further investigated. Also, it is pretty interesting that Nexavant can be applied to the terminal stage of the cancer animal model with a dose increment (Figure 4C).

Current clinical practice’s most promising immune checkpoint inhibitors are anti-PD-1 antibodies^29, 30^. PD-1 antagonists work by disrupting the T cell immune evasion mechanisms employed by cancer cells, thereby enhancing the anti-cancer effect. However, it has been reported that the effectiveness of immune checkpoint inhibitors, such as anti-PD-1 antibodies, is notably diminished in “cold” tumors characterized by a scarcity of immune cells within the tumor microenvironment31, 32. When the anti-PD-1 antibody and Nexavant are co-administered intratumorally, diverse immune cells are infiltrated into the tumor (Figure 5B) and shown to have a synergistic anti-cancer effect (Figure 5). Nexavant can synergistically enhance the recruitment of immune cells to cold tumors with anti-PD-1 antibodies.

In this study, we observed that the anti-cancer efficacies of anti-PD-1 antibodies varied depending on the cancer animal model (partially effective in EO771 and B16F10; less effective in K1735 and EMT6 models). However, the anti-cancer efficacy of Nexavant was more consistent across the test animal models. When administered intratumorally, Nexavant showed a consistent anti-cancer and further additive/synergistic effect with anti-PD-1 antibody in most subcutaneous tumor models. Combination therapies of Nexavant and anti-PD-1 antibody were additive/synergistic in most models. This result suggests that Nexavant can be applicable as a monotherapy or combination therapy, overcoming the limited clinical efficacy of anti-PD-1 antibodies. To expand the idea to a more clinically relevant application, a human anti-PD-1 antibody, Nivolumab, was tested in the transgenic mice expressing a chimeric PD-1 with the extracellular domain of human PD-1. The partial anticancer effect of Nexavant and Nivolumab was synergistic in combinational therapy. Interestingly, the dual therapy was as effective as 83% of complete response in a TNBC cancer model.

The intranasal administration of Nexavant demonstrated a robust anti-cancer activity against lung metastases. The increase of immune cells in the lungs following Nexavant treatment is believed to underlie the anti-cancer effect with recruit groups of immune cells, which can directly induce changes in the tumor microenvironment (Figure 8A). This assay also indicates the better potency of Nexavant compared to Poly(I:C). The study demonstrated that Nexavant should be administered for optimal effectiveness during the early to mid-stage cancer progression. Unlike an intratumoral application, the intranasal combination therapy with anti-PD-1 antibodies did not show improvement. There would be several reasons, such as the anatomic features of the lung as well as the distribution of anti-PD-1 to the lung.

**Figure 8.**
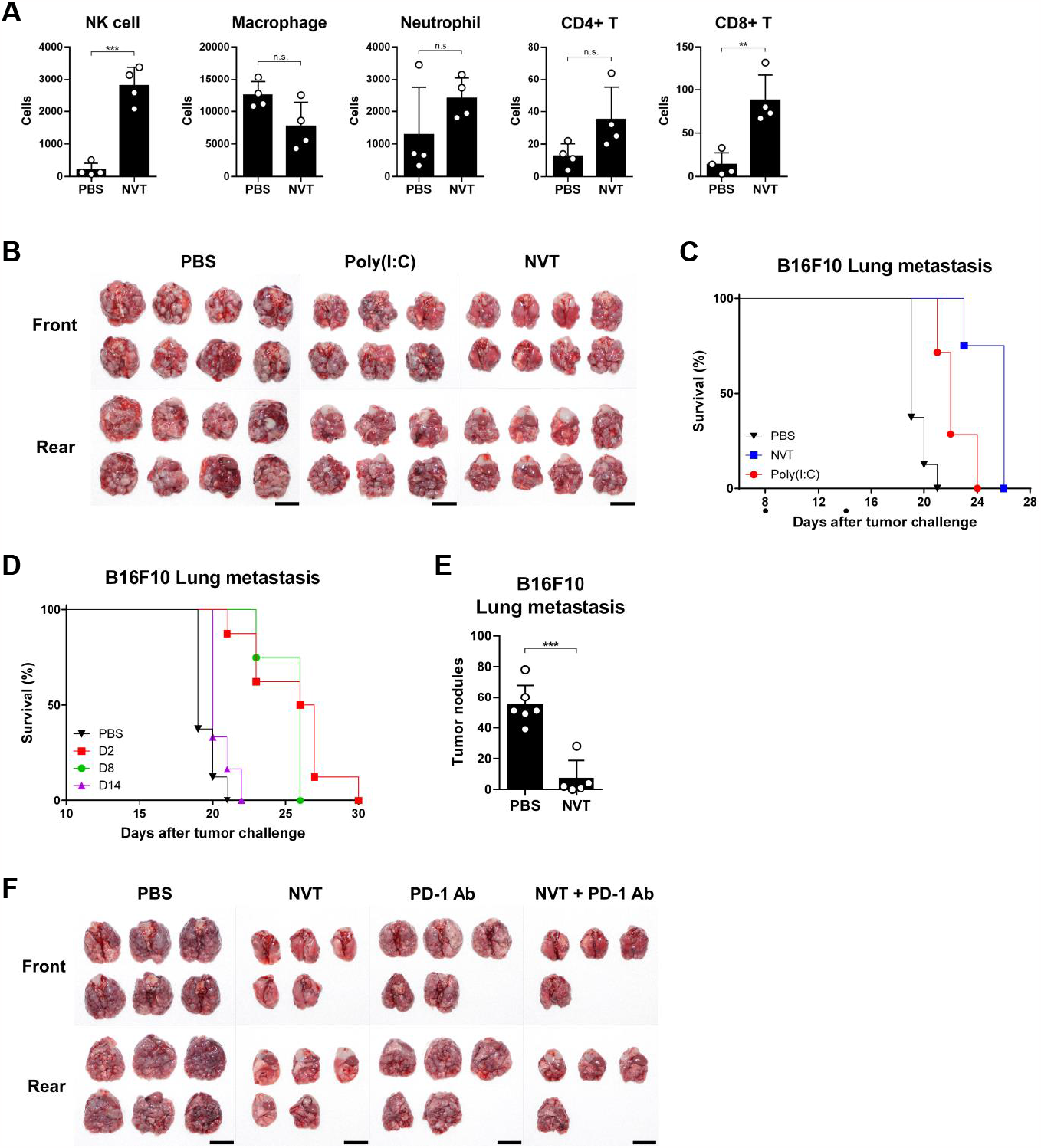
Intranasal administration of Nexavant has a potent antitumor effect, but there is no additive effect when combined with anti-PD-1 antibody. (A) 25 ug of Nexavant was slowly instilled through the nostrils. After 24 hours, bronchoalveolar lavage fluid (BALF) was collected and analyzed by flow cytometry. (B, C) C57bl/6 mice were injected with 5 x 105 B16F10 cells via the tail vein. 25 ug of Nexavant or Poly(I:C) was slowly instilled through the nostrils continuously every two days starting on day 8 or 14. Tumor-bearing lungs (B) were harvested on day 18, and overall survival (C) was monitored. (D, E) C57bl/6 mice were injected with 5 x 105 B16F10 cells via the tail vein. Then, 25 ugs of Nexavant was slowly instilled through the nostrils continuously every two days starting on days 2, 8, or 14, respectively. (E) On day 15, lungs were harvested from B16F10 lung metastases, and tumor nodules were counted. (F) C57bl/6 mice were injected with 5 x 105 B16F10 cells via the tail vein. Then 25 ug Nexavant was instilled via the nostrils, and 200 ug anti-PD-1 antibody was injected intraperitoneally on days 2, 4, 6, 8, and 10. On day 18, B16F10 lung metastasis mice were sacrificed, and lungs were harvested. The gross photograph of a lung with tumor burden was taken at a certain distance.

## 5. Conclusions

In this study, we demonstrated that Nexavant is a distinct TLR3 agonist, which possesses better physicochemical properties (i.e., a feasible quantitation, qualification, thermostability, and more resistance to RNase A for enhanced charters for mass manufacturing and clinical use compared with Poly(I:C). Unlikely Poly(I:C) HMW form, targeting all three dsRNA sensors of TLR3, RIG-I, and MDA5 in both the naked and cationic lipid complex forms, Nexavant targets TLR3 only in the naked form and TLR3 and RIG-I, without MDA5 in the cationic lipid complex form. In situ intratumoral application, Nexavant itself, shows a potent anticancer reagent in diverse cancer animal models by recruiting immune cells to tumor tissue. The synergistic effect of it with anti-PD1 antibody was demonstrated in various cancer models. With the combination of Nivolumab with a chimeric mouse with a human PD-1 domain, the effect also synergizes and suppresses tumor growth by recruiting diverse immune cells into tumors, especially in a cold tumor model. Nexavant shows a potent anticancer effect on the metastatic lung cancer model through an intranasal application. Unlike the intratumoral model, the synergistic effect of the Nexavant/anti-PD-1 combination was not observed in the lung metastasis model.

## Funding

This research was supported by a grant from the Korea Health Technology R&D Project through the Korea Health Industry Development Institute (KHIDI), funded by the Ministry of Health & Welfare, Republic of Korea (grant number: HV22C010200, HV22C010000).

## Conflicts of Funding

S.-H. Lee, Y.-H. Choi, S.M. Kang, M.-G. Lee, S.-B. Hong and D.-H. Kim is a full-time employee of NA Vaccine Institute. A. Debin and E. Perousel are full-time employees of InvivoGen. D.-H. Kim received research grants from the Korea Health Industry Development Institute (KHIDI) and is the inventor of the patent related to this work (patent #: KR102490068B1).

